# Systems Biology Theory Clarification of a Controversy in Pancreatic Beta Cell Regeneration

**DOI:** 10.1101/469320

**Authors:** Haoran Cai, Runtan Cheng, Ruoshi Yuan, Xiaomei Zhu, Ping Ao

## Abstract

Whether new pancreatic beta-cells arise via pre-existing beta-cells or from differentiation of precursor cells − a question of fundamental importance for diabetic therapy − has long been debated. Recent experiments suggest that multipotent precursors from adult mouse pancreas, that give rise to beta-cells, do exist. However, such a finding is at odds with prior evidence that beta-cell expansions occurs exclusively through self-replication. Here we show that these two observations can be partially compatible. We use a systems biology approach to analyze the dynamics of the endogenous molecular-cellular network in the pancreas. Our results show that self-replicating ‘beta-cells’ can themselves be multipotent precursors. In addition, our model predicts heterogeneity in beta-cell regeneration and suggests various differentiation paths of precursors. This work therefore provides a means of reconciling an apparent contradiction in the field, but also sheds light on possible paths of beta-cell regeneration from a systems biology perspective.

## Introduction

Diabetes has brought great public health burden as well as economic costs[1, 2], which is caused by the body’s lack of proper response to insulin production (Insulin resistance) or the pancreas’s inability to produce enough insulin. Both β-cell dysfunction and decreased β-cell mass account for insulin deficiency. It is now recognized that beta-cell loss is a common theme of type 1 diabetes and type 2 diabetes. For example, in patients with type 2 diabetes, the beta-cell mass was found to be reduced by 50%[3]. Thus, many studies have tried to elucidate the mechanisms that control beta cell formation and replacement in order to design regenerative therapy of diabetes[4, 5].

Yet, one highly controversial issue remains unsettled on the postnatal origins of beta cells: whether new pancreatic beta-cells arise via pre-existing beta-cells or differentiation of precursor cells. It has been found that adult beta-cell retains a small capacity for proliferation[6–8]. Surprisingly, a seminal lineage-tracing study found that the fraction of labeled beta-cells remained unchanged over a one-year chase period, suggesting that beta-cell expansion was driven by self-replication without any contribution of precursor cell differentiation[9], which can be further amplified by subsequent studies[10, 11]. On the other hand, several studies have supported the idea that the formation of new endocrine cells is from the pancreatic stem cells since the late 19^th^-century [12–14], Recently, several studies have found that multipotent precursors do exist. A report argued that the beta-cells labeled in the lineage-tracing study may not necessarily be mature beta-cell and can be insulin-expressing precursors that give rise to endocrine cell types[15]. Another group also identified non-insulin-expressing cells in islets that could give rise to new insulin+ cells as a slow renewal for beta-cells.

Here we used a systems biology method, the endogenous molecular-cellular network theory, to clarify the contrasting observations from a systematic view. The theory is intended to analyze the complex biological process via the dynamics of the network system based on the fundamental properties of the biological system[16, 17]. The endogenous molecular-cellular network is composed of essential modules (specified by a set of nodes representing key proteins or signaling pathway) and crosstalk between these modules, and then we analyze the dynamics on the network. We assume cell phenotypes are endogenous attractors underlying the endogenous molecular–cellular network. One of the most prominent predictions of endogenous network theory is the existence of quantitative functional landscape, where locally stable states are interconnected with each other through the transition states. The natural and qualitative consequence of the mathematical setup implies that natural conversion between two cell states may have patterns[17], and the topological structure of the functional landscape can be a natural framework for beta-cell regeneration. We sought to analyze the natural mechanisms at a network level by which beta cells are formed, where the observations of beta-cell expansion can be integrated into a single model.

## Methods

### Construction of an endogenous network of pancreas development

In this work, we chose a set of essential proteins to depict pancreatic core regulatory structures (detailed description in the unpublished paper). These proteins and their causal interactions form the core endogenous network of the pancreas.

### Quantitative description and analysis

A set of ordinary differential equations (ODEs) were obtained to quantify the core dynamics on the pancreatic endogenous network. The dynamics of the activation/expression level of each protein x is governed by

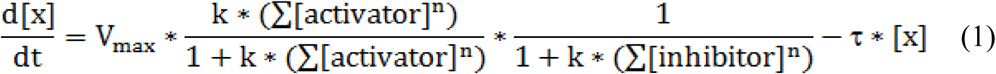

where *V*_*max*_ represents the maximal production rate of protein *x, n* represents Hill coefficients, and *k* represents dissociation constant. Specifically, the relative expression level of each protein was normalized to range from 0 to 1. The maximal production rate *V*_*max*_ and degradation rate **τ** were taken as 1. Here the values of *n* and *k* were 3 and 10 while we conduct multiple simulation varying n and k within a reasonable range to grasp the key feature of activation or inhibition. The threshold of the sigmoid-shaped function, at which the value of *x* was expected to be half maximal. Two independent algorithms, random sampling and Newton’s method, were adopted to calculate the robust fixed points of the dynamical system (see Supplementary Materials).

In the dynamic system (Eqn. 1), we perturbed the system with small random noise when it stayed at an unstable state (transition state or hyper-transition state) to obtain the topological structure of landscape. We utilized random-perturbed states as the initial value of the system and let the dynamical systems iterate at the constraints and tracked routes of system evolution. We obtained the trajectories from each unstable state to it connected stable states and recorded the unstable states that each trajectory passed through.

Independent datasets[18] are used to validate the model results. Firstly, four pathways expression levels are denoted by the average of targeted proteins (pathways). We averaged expression level by cell type annotation. Then Z-score normalization is conducted respectively for single cell transcriptome and modeling results (12 stable states and 23 transition states) over each protein (pathways). Eventually, we do linearly rescale for all the expression value to 0 − 1.

## Results

We obtained 12 stable states and 23 transition states in the simulations of our model (see in Table S2, Table S3). Each state is depicted by the combinational expression level of endogenous network nodes. We assumed pancreatic phenotypes are robust stable states of the endogenous network, thus biological meanings were linked to the stable states emerging from the endogenous network (see in Figure 2, Figure S2). The interconnections among phenotypic states reveal the lineage conversion routes in the pancreas, making it possible to understand the maintenance of pancreas homeostasis from a systematic view. In our work, a beta cell proliferation landscape could be obtained describing how states connect with each other, part of the whole pancreatic landscape (Figure 3).

**Figure 1.**
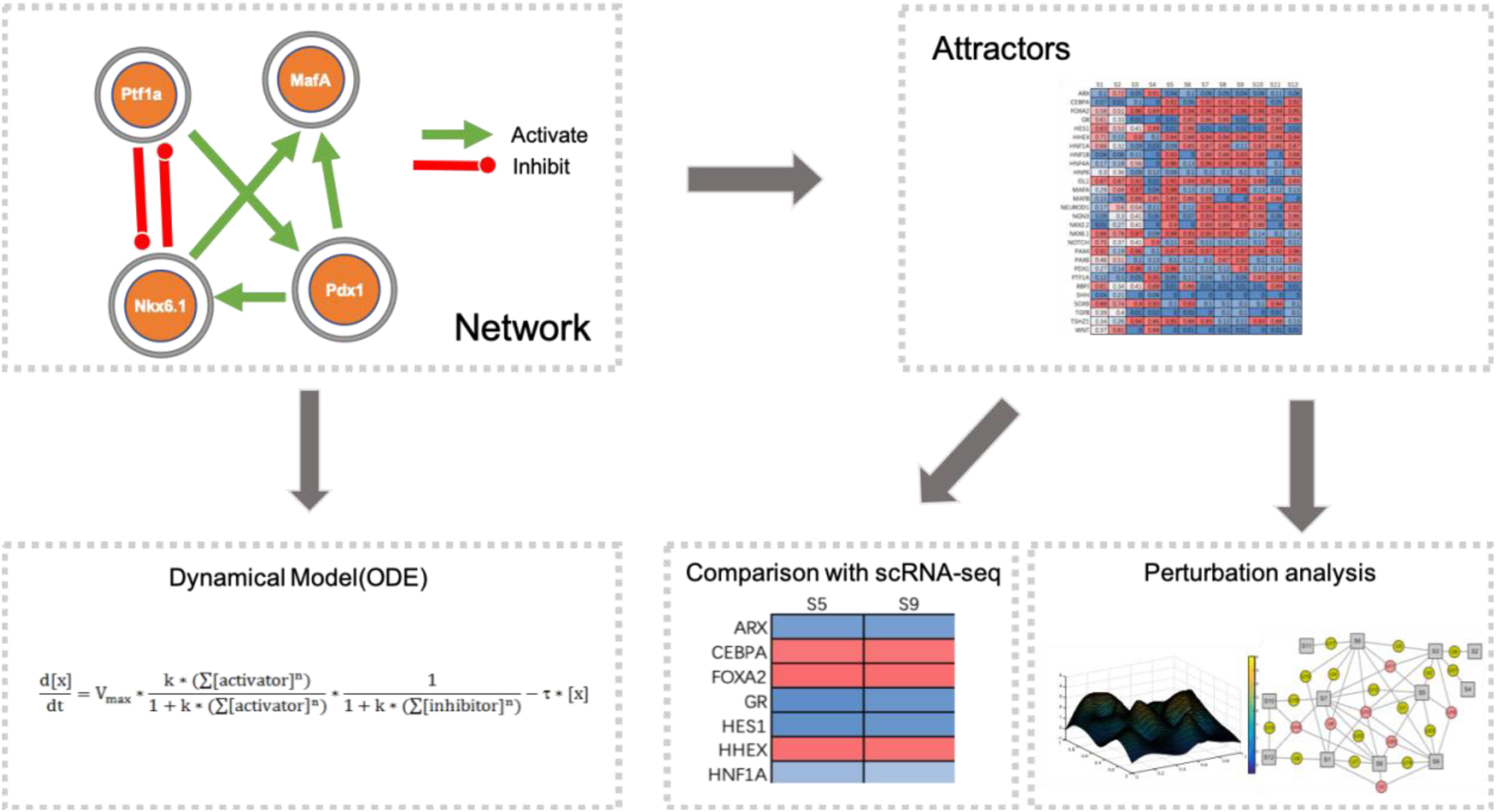
Schematics of endogenous molecular-cellular network modelling. The interactions were collected from the literature. Ordinary differential equations (described in Methods and Supplementary Material) were used to compute the attractors generated by the constructed pancreatic developmental network structure. Two algorithms (see Supplementary Materials) were performed, demonstrating robustness of the simulation results. Comparison of gene expression levels predicted by the attractors with single cell RNA-seq data validated our scientific simulation results. Multiple phenotypes within pancreas corresponded to the attractors of network dynamics.

**Figure 2.**
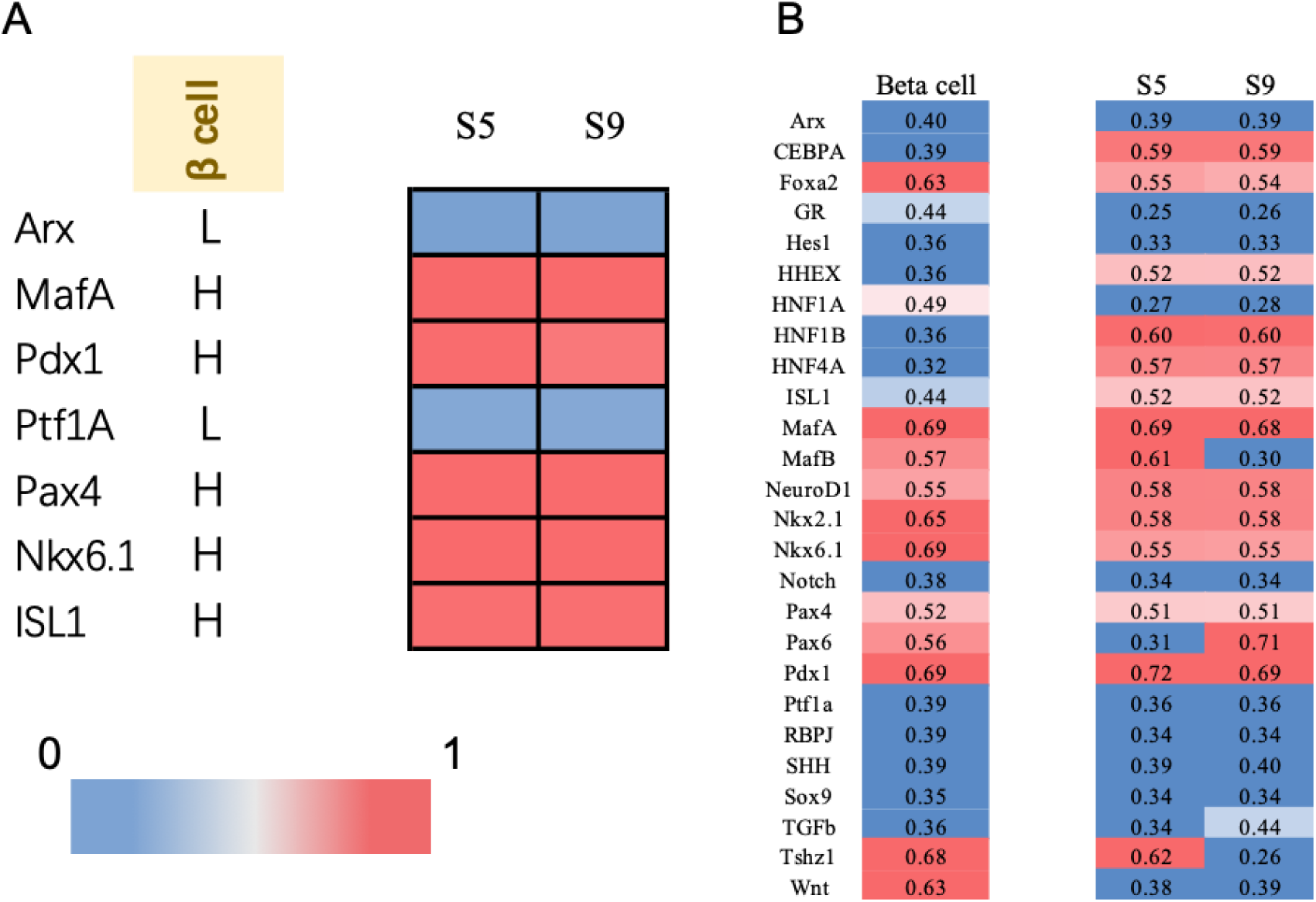
Linking biological meaning to stable states and its validation comparing with RNA-Seq. A. We used the known markers (L denotes Low expression, H denotes High expression), we could easily link the cell type to corresponding stable states generated from the endogenous network (See Figure S2). **B** The biological meanings of beta cell states were validated at the molecular level. We selected the relevant expression data in a published dataset[18] and set a threshold to find out the high or low expressed status of each gene. When we set the threshold as 0.5, the agreement ratio was 71.2%

**Figure 3.**
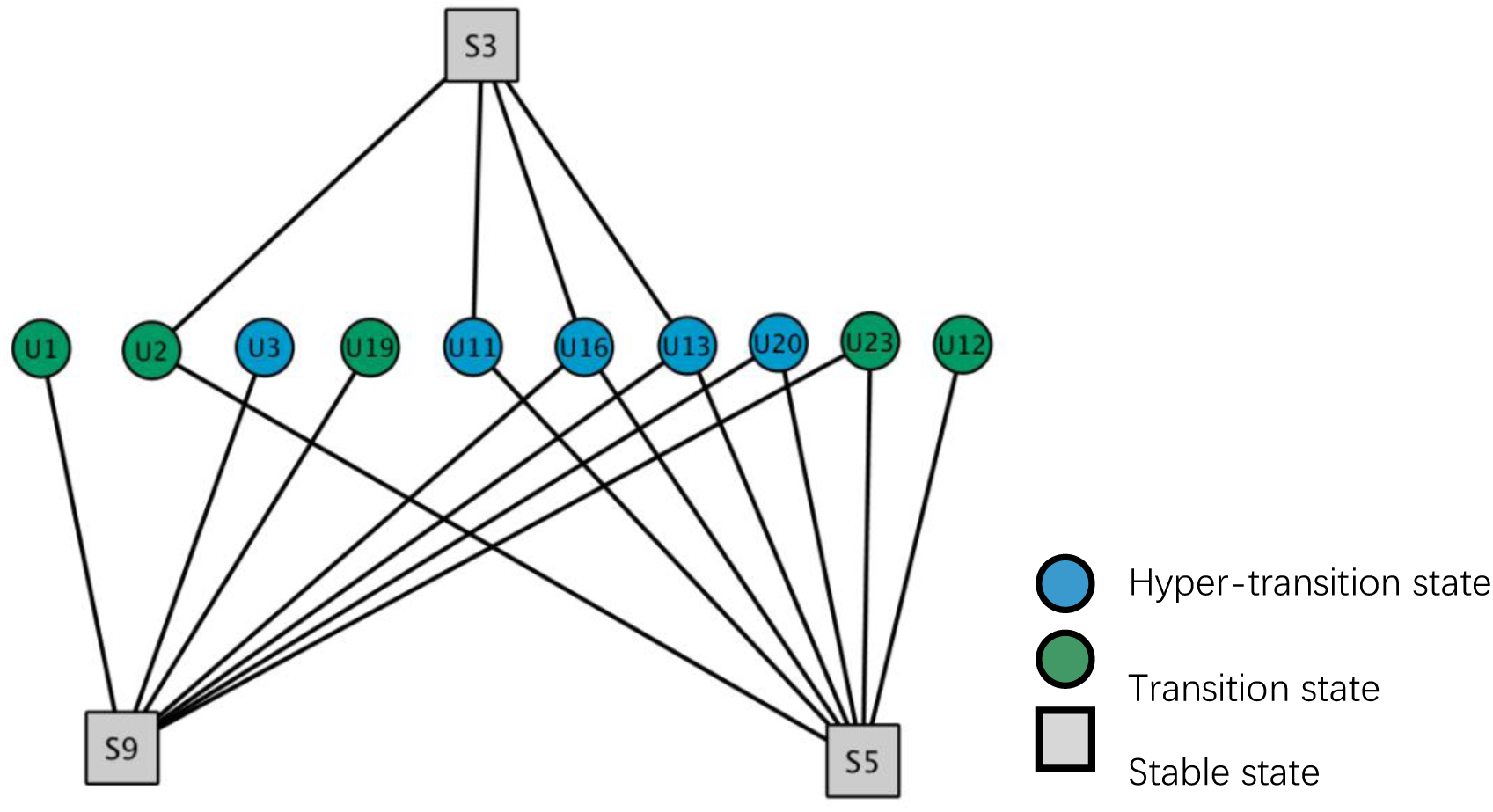
The network dynamics incorporated the seemingly conflicted beta-cell expansion models. This state connection graph is a part of the whole landscape (See Table S1) through perturbation analysis. Each state is depicted by the expression level of a set of molecular we selected to form the endogenous network. S5, S9 represented beta cell states while all the other phenotypic states including stable states S3 and transition state/hyper-transition state retain the potential to differentiate into the beta cell. Stable state: all the eigenvalues of the Jacobian matrix of dynamical system at this state were negative; Transition state: one eigenvalue of the Jacobian matrix at this state was positive while the others were negative; Hyper-transition state: more than one positive eigenvalues of the Jacobian matrix at this state were positive.

By state-connected graph, multiple potential sources including 1 stable state and 11 transition states for expansion can be identified (Figure 3). By examining the expression pattern, U23, S3, U2 states are found to be insulin-producing cells that could give rise to formation of new beta-cells and other phenotypic of states (See in Figure S1) according to our modeling results, as markers that activate Ins are highly expressed at these states (Figure 4). This can be validated by the report that identified an multipotent insulin-expressing population as the precursor of beta-cells[15]. While states such as U11, U12, U20 that do not express a high level of Pdx1, MafA (Figure 4) are insulin-negative states in our model. And they maintained the capacity to give rise to new beta-cells as well (Figure 3), consistent with the postulations from [19] that Insulin-negative precursors participate in the renewal of the beta-cell mass during aging. In this way our model not only explained two evidences straightforwardly but integrated the evidences of existence of precursor cells into a single framework from a systematic view.

**Figure 4.**
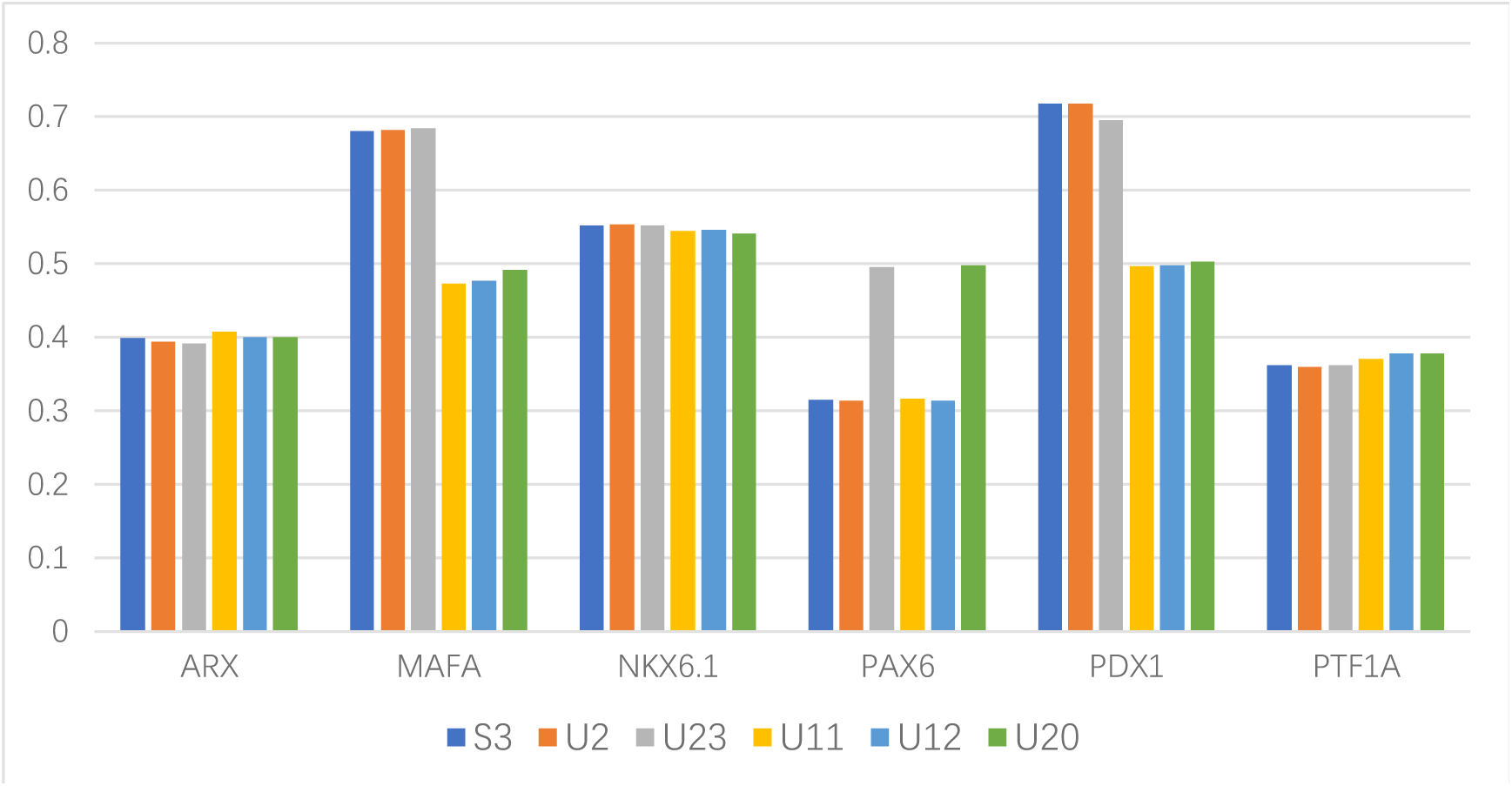
Modeling results characterized the insulin expression features of putative beta cell precursor through a quantitative expression level of proteins. States that highly express MafA, Nkx6.1, Pax6, Pdx1 and low express Arx, Ptf1a are more likely to express insulin[28–31]. State S3, U2, U23 were assumed to be insulin-positive. Due to the lack of expression of MafA, Pdx1, U11, U12, U20 are probably insulin-negative cells.

Furthermore, our model suggested that the sources of beta-cells regeneration may even be more heterogeneous than expected. For example, some of the precursor states are unipotent to beta-cells according to the complete landscape such as U2, U16, U23, suggesting a capacity in vivo that might be exploited to exclusively generating beta-cells for therapeutic cell replacement. Furthermore, as the newly differentiated beta-cells are hard to be distinguished from pre-existing population in many experimental models, the expression characteristics of multiple possible pre-beta states can be obtained (see in Table S2, Table S3) through our modeling to enable future experimental validations for post-natal origin of beta-cells.

## Discussion

Whether new beta cells arise from differentiation or by the proliferation of existing beta cells has remained a highly controversial issue for decades. Dor’s study which used an mice model to trace the fraction of insulin-expressing cells suggested that self replication was the major source of beta-cell regeneration exclusively[9]. The existence of progenitor-like insulin-positive states in our model, distinct from mature beta-cells, was probably the partial reason why they found the new insulin-positive cells are differentiated from existing insulin-expressing cells. Because the precursor cells can express insulin, lineage labelling of beta cells by the insulin expression could not discriminate between whether it is self-replication of pre-existing mature beta cells or differentiation from stem cells that give rise to the formation of new beta-cells.

However, two evidences demonstrated that the multipotent insulin expressing precursors could not provide total reconciliation with the work of [9]. First, insulin-negative states are possible to contributed to new insulin+ cells replacement both theoretically and experimentally. Secondly, the precursor beta cells identified before are multipotent[15] and there is no unipotent insulin expressing state in our landscape while Dor and his colleagues found the labeled cells only give rise to beta cells. This could be attributed by techniques: One reason related is that the staining method that Dor took probably hindered the visualization of labeled cells giving rise to states other than beta cells. In addition, it may overlook the cases that the pre-existing mature beta cells retain the capacity to dedifferentiate to an unipotent progenitor state and re-differentiate back, which indeed has been reported by several studies[20, 21]. Since these precursor cells are quite similar with the mature beta cells, it might be hard to discriminate them in experimental models. Another reason might concern with the small fraction of progenitor states: the lineage tracing results merely reflect the behaviors of mature beta-cells.

Enlightened by pancreatic development endogenous network, we constructed the quantitative landscape of pancreatic development without prior knowledge of beta-cell expansion. Emerged stable states are linked to phenotypic states within the pancreas, reproducing the core features of phenotypic states within pancreas to better understand the beta cell replacement. We conducted perturbation analysis to generate topological graph describing interconnection among states, serving as the guideline to understand the beta-cell replacement. The roadmap of beta cell regeneration can be established: Various possible states retain the capacity to give rise to new beta cells either during development, aging or even under injury. Our results supported that the new post-natal beta cells can originate from the unmatured pre-beta cells. The precursors can be rather heterogenous, characterized by combinational expression level of genes(proteins) in our landscape. Besides, we showed that pre-existing beta-cells could transiently dedifferentiate to a progenitor-like state and facilitate the beta-cell replacement, which can be the case hindered by the experimental techniques. The observations were integrated into a single model, and an explanation of the beta-cell origins within adult pancreas has been obtained from a systems biology theory. One remaining question is that to which extent the beta-cell expanded by self-replication or from precursor cells. It is also of interests to know conditions that a certain phenotypic state of cells will occur and contribute more to the beta-cell expansion. We acknowledge that the network has been greatly simplified. It is expected that a more comprehensive network can reproduce more detailed features through the inclusion of more modules, for example, cell cycle. These issues require an explicit inclusion of stochastic effects, where the potential energy landscape can be used to explore more detailed issues[22, 23].

Apart from elucidating the controversy in this work, the dynamical network system we built may have other applications: Our model does not exclude the possibility that other terminally differentiated phenotypes of cells trans-differentiate into beta-cells for their expansion. In the light of our hypothesis, these trans-differentiation behaviors could correspond to the states traveling in the landscape (Figure 3) as well. Indeed, stem cells have been observed in multiple experiment models[24–26]. Hence, it is possible that other multiple routes can generate new beta-cells. Our model provides a framework to understand the interconversion of cell states during aging or embryonic development. The patterns and preferred routes that our model implies can be further studied to predict potential target genes and develop successful therapies for beta cell regeneration in the treatment of diabetes.

## Data availability

The published data sets used in this manuscript are available through the following accession numbers: SMART-seq2 platform pancreas data by Segerstolpe *et al.* [18], ArrayExpres<u>s E-MTAB-5061.</u>

