## Supplemental Tables and Figures for "Systems Biology Theory Clarification of a Controversy in Pancreatic Beta Cell Regeneration"

#### The construction of the core endogenous network of pancreatic cell-fate determination process

Under the theoretical framework of endogenous networks, development and functional execution are regulated by the same core endogenous network. In the process of pancreas development, cell differentiation and fate determination are strictly regulated. Although many genetic and epigenetic regulators are involved in development, it has been found in past studies that the expression of a few lineage-specific transcription factors are the master regulator and play determined roles[1–3]. For example, deletion of Ptf1a leads to redistribution of pancreatic precursor progenitor subpopulations to other endoderm lineages; alpha cell determinant Arx and beta cell determinant Pax4, cross-inhibition between Nkx6.1 and Pdx1 separates the two lineage[1]. In addition, due to the complexity of biological systems, we do not integrate all the gene regulatory factors, but generalize the entire system through the activities of some key proteins related to the fate of pancreatic cells. The pancreatic developmental network (Table S1) can be obtained by reviewing the literature to determine the interaction between factors one by one. It should be pointed out the results of low-throughput basic molecular experiments and biochemical experiments were selected, and high-throughput and bioinformatics analysis results were not taken. For example, a study found that NKX6.1 and NKX6.2 can directly inhibit the expression of Ptf1a gene[4], which can obtain the inhibition relationship between network nodes. These factors are mutually regulated by direct or indirect activation inhibition relationships, forming a set of conserved, closed, endogenous networks that perform specific functions. In our study, we also conducted validation of the agents as to whether they are strong enough to account for the whole system since our network just select 28 agents to describe it. The clustering algorithm[5] were conducted using the two different inputs genes(the whole genome and the selected agents) The results showed a great consistency between these two inputs (Results not shown here).

#### Quantitative model formation of the endogenous network

A set of ordinary differential equations (ODEs) were obtained to quantify the core dynamics on the pancreatic endogenous network. The dynamics of the activation/expression level of each protein x is governed by

$$\frac{d[x]}{dt} = V_{max} * \frac{k * (\sum[\text{activator}]^n)}{1 + k * (\sum[\text{activator}]^n)} * \frac{1}{1 + k * (\sum[\text{inhibitor}]^n)} - \tau * [x] \quad (1)$$

where $V_{max}$ represents the maximal production rate of protein $x$, $n$ represents Hill coefficients, and $k$ represents dissociation constant. Specifically, the relative expression level of each protein was normalized to range from 0 to 1. The maximal production rate $V_{max}$ and degradation rate $\tau$ were taken as 1. Here the values of $n$ and $k$ were 3 and 10 while we conduct multiple simulation varying n and k within a reasonable range to grasp the key feature of activation or inhibition. The threshold of the sigmoid-shaped function, at which the value of $x$ was expected to be half maximal.

#### Algorithms for fixed points calculation in the ODE dynamical system

Algorithm 1

This algorithm generated a random vector $x_0$ as an initial value of the system, then iterated $x_0$ by using $x_{t+1} = x_t + \Delta t \times f(x_t)$. Its convergence is judged by $(x_{t+1} - x_t)^2 < \varepsilon, f(x_t)^2 < \delta$ after a considerable number (N) of iteration. Let $\Delta t = 0.1, N = 800, \varepsilon = 10^{-8}, \delta = 10^{-8}$. If $x_t$ met the judgments, we recorded the $x_t$ as a stable state. By randomly sampling the initial vectors, for example, 10000 times, the stable states can be obtained. If the random samplings were repeated 10000000 or more and we still obtained the same results, then these robust stable states are our primary results.

Algorithm 2

This algorithm is able to calculate both stable states and unstable states. Fixed points could be acquired through solving the nonlinear equation $\frac{dx(t)}{dt} = f(x) = 0$ by using Newton's method. We generated a random vector $x_0$ as an initial state of the dynamical system, then updated $x_0$ by Newton’s method to find the solution $x^*$ which satisfied $f(x^*) = 0$ by using the fsolve function in MATLAB. If all the eigenvalues of the Jacobian matrix of $f(x)$ at $x^*$ were negative, $x_i$ was a stable state. If there was at least an eigenvalue at $x^*$ was positive, then we recorded the solution $x_i$ as an unstable state including transition state with a positive eigenvalue and hyper-transition state with more than one positive eigenvalue. By randomly sampling the initial vectors, for example, 10000 times, the stable states and the unstable states were recorded. If the sampling times were repeated 10000000 or more times and we still obtained the identical results, then we used these stable states and unstable states as primary results.

#### Obtaining topological structure of landscape through State interconnection analysis

In the dynamic system of ODEs, we perturbed the system randomly with small random vectors Δx (|Δx| ≤ 0.005) when it stayed at an unstable state. We utilized random-perturbed states as the initial value of the system and let the dynamical systems iterate at the constraints of and tracked routes of system evolution. To judge whether the trajectories would pass through the other unstable states around, we enumerated all the calculated unstable states and judge the convergence by $(x_t - x_{unstable\ node})^2 < \varepsilon_1$ at each updated step. After a considerable times (N) of iteration, we judged its convergence by $(x_{t+1} - x_t)^2 < \varepsilon, f(x_t)^2 < \delta$ to find out the local stable states that the system finally reached. Here, we let $\Delta t = 0.1, N = 800, \varepsilon_1 = 10^{-3}, \varepsilon = 10^{-8}, \delta = 10^{-8}$. We obtained the trajectories from each unstable state to it connected stable states and recorded the unstable states that each trajectory passed through. The random perturbations were repeated 1000 times, and we recorded the unstable states and stable states along each trajectory. If the perturbations were repeated 10000 or more times and we still obtained the identical results, then we used them as primary results.

#### Validation of the biological meanings with gene profiling data at molecular level

The 12 steady states can be obtained from the network. Each state is depicted by the combinational expression level of endogenous network nodes. They were linked to biological meanings, which are phenotypes. Further, they were compared with those obtained from human mature pancreas RNA-seq data[6] to further verify the network calculation results. Firstly, four pathways expression levels are denoted by the average of targeted proteins (pathways). We averaged expression level by cell type annotation. Then Z-score normalization is conducted respectively for single cell transcriptome and modeling results (12 stable states and 23 transition states) over each protein (pathways). Eventually, we do linearly rescale for all the expression value to 0 – 1. We set the threshold to represent whether each steady state is activated or suppressed. If the threshold is 0.5, the expression above 0.5 is considered to be activated, otherwise it is inhibition. The threshold was set within a reasonable range to decide the On/Off status of each gene to considerably examine the results. The comparison is showed in Figure S3 and the agreement ratio under different threshold see in Table S4

### Supplementary Tables

**Table S1. The causal interactions in the endogenous network of pancreatic development process.** The agents in the first column were activated or inhibited by these molecular-cellular agents in the second and third column respectively.

| Nodes | Activator | Inhibitor |
|---|---|---|
| ARX | TSHZ1; FOXA2; ISL1; NGN3; MAFA | PAX4; NKX2.2; PDX1 |
| CEBPA | HHEX; HNF4A | NOTCH; SHH; SOX9 |
| FOXA2 | SOX9; HNF1B; PDX1; HHEX; HNF6; SHH; CEBPA | TGFB |
| GR | HNF1A; TGFB | HNF6 |
| HES1 | NOTCH; TGFB | |
| HHEX | TGFB; NKX2.2; FOXA2 | ARX |
| HNF1A | HNF6; HHEX | PDX1 |
| HNF1B | HNF6; PDX1; CEBPA | SOX9 |
| HNF4A | HNF1A; HNF1B; PDX1; HHEX; CEBPA | NOTCH |
| HNF6 | SOX9; PTF1A; HNF1B; HNF4A; CEBPA; TGFB | FOXA2 |
| ISL1 | NKX6.1; NEUROD1 | |
| MAFA | ISL1; NGN3; PDX1; NEUROD1; NKX6.1; HNF1A | GR |
| MAFB | TSHZ1 | PAX6 |
| NEUROD1 | NGN3; WNT; NKX2.2; NKX6.1 | NOTCH |
| NGN3 | PDX1; NKX6.1; HNF1B; HNF6; SOX9 | HES1; NOTCH |
| NKX2.2 | NGN3; SHH | |
| NKX6.1 | PAX4; TSHZ1; MAFA; PDX1; NKX2.2 | PTF1A |
| NOTCH | PTF1A; NGN3; HNF1A; HNF1B | CEBPA |
| PAX4 | NGN3; PDX1; ISL1; FOXA2; HNF1A | ARX |
| PAX6 | ISL1; NOTCH; PDX1; ARX | TGFB; TSHZ1 |
| PDX1 | FOXA2; HNF6; TSHZ1; MAFB; PTF1A; RBPJ | GR; ARX |
| PTF1A | SOX9; NOTCH; HNF1B; HES1 | NKX6.1; MAFA |
| RBPJ | NOTCH | |
| SHH | ARX; TGFB | FOXA2; HES1 |
| SOX9 | FOXA2; HES1 | CEBPA |
| TGFB | PAX6 | CEBPA; HNF6 |

| | | |
|---|---|---|
| TSHZ1 | PDX1; NKX2.2; FOXA2; | PAX6 |
| WNT | ARX; TGFB | |

**Table S2. Expression levels of genes in stable states of the core endogenous network**

| | S1 | S2 | S3 | S4 | S5 | S6 | S7 | S8 | S9 | S10 | S11 | S12 |
|---|---|---|---|---|---|---|---|---|---|---|---|---|
| ARX | 0.1 | 0.72 | 0.05 | 0.91 | 0.04 | 0.1 | 0.06 | 0.06 | 0.04 | 0.06 | 0.11 | 0.06 |
| CEBPA | 0.07 | 0.01 | 0.1 | 0 | 0.92 | 0.06 | 0.92 | 0.92 | 0.92 | 0.92 | 0.05 | 0.92 |
| FOXA2 | 0.58 | 0.51 | 0.96 | 0.89 | 0.97 | 0.94 | 0.96 | 0.95 | 0.96 | 0.96 | 0.94 | 0.95 |
| GR | 0.61 | 0.33 | 0.01 | 0 | 0.01 | 0.85 | 0.86 | 0.86 | 0.02 | 0.86 | 0.85 | 0.86 |
| HES1 | 0.83 | 0.53 | 0.41 | 0.88 | 0.01 | 0.86 | 0.01 | 0.02 | 0.02 | 0.01 | 0.89 | 0.02 |
| HHEX | 0.71 | 0.15 | 0.9 | 0.1 | 0.94 | 0.88 | 0.94 | 0.94 | 0.94 | 0.94 | 0.88 | 0.94 |
| HNF1A | 0.66 | 0.32 | 0.09 | 0.03 | 0.09 | 0.85 | 0.87 | 0.88 | 0.11 | 0.87 | 0.85 | 0.87 |
| HNF1B | 0.04 | 0.06 | 0.11 | 0 | 0.93 | 0 | 0.88 | 0.88 | 0.93 | 0.88 | 0 | 0.88 |
| HNF4A | 0.17 | 0.18 | 0.56 | 0 | 0.96 | 0.13 | 0.96 | 0.96 | 0.96 | 0.96 | 0.1 | 0.96 |
| HNF6 | 0.3 | 0.36 | 0.09 | 0.12 | 0.09 | 0.1 | 0.1 | 0.1 | 0.1 | 0.1 | 0.1 | 0.1 |
| ISL1 | 0.87 | 0.87 | 0.92 | 0.02 | 0.95 | 0.89 | 0.95 | 0.94 | 0.95 | 0.89 | 0.01 | 0.89 |
| MAFA | 0.29 | 0.68 | 0.97 | 0.04 | 0.98 | 0.13 | 0.13 | 0.13 | 0.98 | 0.13 | 0.12 | 0.13 |
| MAFB | 0.15 | 0.06 | 0.88 | 0.85 | 0.89 | 0.86 | 0.88 | 0 | 0 | 0.88 | 0.86 | 0 |
| NEUROD1 | 0.17 | 0.6 | 0.54 | 0.11 | 0.95 | 0.12 | 0.95 | 0.95 | 0.95 | 0.92 | 0 | 0.92 |
| NGN3 | 0.09 | 0.3 | 0.41 | 0.06 | 0.95 | 0.07 | 0.93 | 0.93 | 0.95 | 0.86 | 0.06 | 0.86 |
| NKX2.2 | 0.01 | 0.27 | 0.41 | 0 | 0.9 | 0 | 0.89 | 0.89 | 0.9 | 0.86 | 0 | 0.86 |
| NKX6.1 | 0.88 | 0.78 | 0.97 | 0.09 | 0.98 | 0.93 | 0.95 | 0.93 | 0.97 | 0.14 | 0.1 | 0.14 |
| NOTCH | 0.75 | 0.37 | 0.41 | 0.9 | 0.11 | 0.86 | 0.11 | 0.11 | 0.11 | 0.11 | 0.93 | 0.11 |
| PAX4 | 0.91 | 0.19 | 0.96 | 0.1 | 0.97 | 0.95 | 0.97 | 0.97 | 0.97 | 0.96 | 0.92 | 0.96 |
| PAX6 | 0.46 | 0.51 | 0.1 | 0.13 | 0.1 | 0.12 | 0.1 | 0.87 | 0.92 | 0.1 | 0.11 | 0.85 |
| PDX1 | 0.27 | 0.14 | 0.96 | 0.12 | 0.96 | 0.13 | 0.13 | 0.12 | 0.9 | 0.13 | 0.14 | 0.13 |
| PTF1A | 0.12 | 0.1 | 0.05 | 0.95 | 0.05 | 0.11 | 0.09 | 0.1 | 0.05 | 0.83 | 0.93 | 0.83 |
| RBPJ | 0.81 | 0.34 | 0.41 | 0.88 | 0.01 | 0.86 | 0.01 | 0.01 | 0.01 | 0.01 | 0.89 | 0.01 |
| SHH | 0.04 | 0.21 | 0 | 0.06 | 0 | 0 | 0 | 0 | 0 | 0 | 0 | 0 |
| SOX9 | 0.88 | 0.74 | 0.9 | 0.93 | 0.1 | 0.93 | 0.1 | 0.1 | 0.1 | 0.1 | 0.94 | 0.1 |
| TGFB | 0.39 | 0.4 | 0.01 | 0.02 | 0 | 0.02 | 0 | 0.1 | 0.1 | 0 | 0.01 | 0.1 |
| TSHZ1 | 0.34 | 0.26 | 0.94 | 0.86 | 0.95 | 0.88 | 0.93 | 0.12 | 0.11 | 0.93 | 0.88 | 0.13 |
| WNT | 0.37 | 0.81 | 0 | 0.88 | 0 | 0.01 | 0 | 0.01 | 0.01 | 0 | 0.01 | 0.01 |

**Table S3. Transition states of the pancreatic endogenous network.**

| | U1 | U2 | U3 | U4 | U5 | U6 | U7 | U8 | U9 | U10 | U11 | U12 | U13 | U14 | U15 | U16 | U17 | U18 | U19 | U20 | U21 | U22 | U23 |
|---|---|---|---|---|---|---|---|---|---|---|---|---|---|---|---|---|---|---|---|---|---|---|---|
| ARX | 0.07 | 0.04 | 0.08 | 0.07 | 0.09 | 0.54 | 0.08 | 0.06 | 0.07 | 0.1 | 0.07 | 0.05 | 0.07 | 0.06 | 0.06 | 0.05 | 0.1 | 0.06 | 0.05 | 0.05 | 0.91 | 0.06 | 0.04 |
| CEBPA | 0.51 | 0.56 | 0.5 | 0.57 | 0.09 | 0.05 | 0.56 | 0.92 | 0.57 | 0.06 | 0.55 | 0.92 | 0.54 | 0.92 | 0.92 | 0.55 | 0.05 | 0.92 | 0.92 | 0.92 | 0 | 0.92 | 0.92 |
| FOXA2 | 0.73 | 0.96 | 0.71 | 0.92 | 0.94 | 0.52 | 0.75 | 0.95 | 0.86 | 0.64 | 0.92 | 0.96 | 0.85 | 0.96 | 0.96 | 0.88 | 0.94 | 0.96 | 0.95 | 0.96 | 0.89 | 0.96 | 0.97 |
| GR | 0.24 | 0.01 | 0.35 | 0.86 | 0.48 | 0.34 | 0.82 | 0.86 | 0.85 | 0.73 | 0.48 | 0.48 | 0.43 | 0.86 | 0.86 | 0.09 | 0.85 | 0.86 | 0.45 | 0.45 | 0 | 0.86 | 0.01 |
| HES1 | 0.38 | 0.23 | 0.39 | 0.24 | 0.57 | 0.53 | 0.36 | 0.02 | 0.28 | 0.85 | 0.23 | 0.01 | 0.29 | 0.01 | 0.02 | 0.28 | 0.87 | 0.01 | 0.02 | 0.01 | 0.76 | 0.01 | 0.01 |
| HHEX | 0.85 | 0.93 | 0.84 | 0.92 | 0.88 | 0.27 | 0.85 | 0.94 | 0.9 | 0.75 | 0.92 | 0.94 | 0.9 | 0.94 | 0.94 | 0.91 | 0.88 | 0.94 | 0.94 | 0.94 | 0.1 | 0.94 | 0.94 |
| HNF1A | 0.19 | 0.09 | 0.32 | 0.87 | 0.46 | 0.32 | 0.85 | 0.88 | 0.86 | 0.76 | 0.45 | 0.45 | 0.41 | 0.88 | 0.87 | 0.11 | 0.85 | 0.87 | 0.43 | 0.44 | 0.02 | 0.88 | 0.1 |
| HNF1B | 0.59 | 0.68 | 0.52 | 0.51 | 0.06 | 0.07 | 0.51 | 0.88 | 0.51 | 0.02 | 0.53 | 0.89 | 0.53 | 0.88 | 0.88 | 0.66 | 0 | 0.88 | 0.89 | 0.89 | 0 | 0.88 | 0.93 |
| HNF4A | 0.69 | 0.73 | 0.68 | 0.72 | 0.38 | 0.26 | 0.7 | 0.96 | 0.71 | 0.14 | 0.71 | 0.95 | 0.7 | 0.96 | 0.96 | 0.72 | 0.12 | 0.96 | 0.95 | 0.95 | 0.01 | 0.96 | 0.96 |
| HNF6 | 0.18 | 0.09 | 0.19 | 0.1 | 0.1 | 0.34 | 0.17 | 0.1 | 0.12 | 0.24 | 0.1 | 0.1 | 0.12 | 0.1 | 0.1 | 0.11 | 0.1 | 0.1 | 0.1 | 0.1 | 0.11 | 0.1 | 0.1 |
| ISL1 | 0.92 | 0.93 | 0.92 | 0.92 | 0.9 | 0.87 | 0.91 | 0.9 | 0.92 | 0.87 | 0.93 | 0.95 | 0.92 | 0.9 | 0.89 | 0.93 | 0.51 | 0.89 | 0.94 | 0.94 | 0.24 | 0.94 | 0.95 |
| MAFA | 0.84 | 0.97 | 0.66 | 0.13 | 0.45 | 0.66 | 0.15 | 0.13 | 0.13 | 0.2 | 0.46 | 0.47 | 0.53 | 0.13 | 0.13 | 0.96 | 0.12 | 0.13 | 0.51 | 0.5 | 0.33 | 0.13 | 0.98 |
| MAFB | 0.03 | 0.89 | 0.04 | 0.87 | 0.86 | 0.09 | 0.02 | 0 | 0.21 | 0.28 | 0.88 | 0.88 | 0.2 | 0.2 | 0.2 | 0.21 | 0.86 | 0.88 | 0 | 0.22 | 0.85 | 0.21 | 0.24 |
| NEUROD1 | 0.68 | 0.72 | 0.68 | 0.71 | 0.38 | 0.58 | 0.68 | 0.92 | 0.7 | 0.14 | 0.72 | 0.95 | 0.71 | 0.92 | 0.92 | 0.71 | 0.07 | 0.92 | 0.95 | 0.95 | 0.21 | 0.95 | 0.95 |
| NGN3 | 0.49 | 0.67 | 0.48 | 0.63 | 0.22 | 0.3 | 0.5 | 0.87 | 0.59 | 0.07 | 0.65 | 0.93 | 0.59 | 0.87 | 0.86 | 0.62 | 0.06 | 0.87 | 0.93 | 0.93 | 0.11 | 0.93 | 0.95 |
| NKX2.2 | 0.54 | 0.75 | 0.53 | 0.71 | 0.1 | 0.24 | 0.55 | 0.87 | 0.67 | 0 | 0.73 | 0.89 | 0.67 | 0.87 | 0.86 | 0.7 | 0 | 0.87 | 0.89 | 0.89 | 0.02 | 0.89 | 0.9 |
| NKX6.1 | 0.95 | 0.97 | 0.93 | 0.95 | 0.94 | 0.79 | 0.91 | 0.42 | 0.92 | 0.88 | 0.95 | 0.96 | 0.93 | 0.41 | 0.14 | 0.97 | 0.47 | 0.41 | 0.94 | 0.95 | 0.28 | 0.94 | 0.97 |
| NOTCH | 0.33 | 0.31 | 0.32 | 0.32 | 0.51 | 0.38 | 0.33 | 0.11 | 0.32 | 0.82 | 0.31 | 0.11 | 0.32 | 0.11 | 0.11 | 0.32 | 0.88 | 0.11 | 0.11 | 0.11 | 0.68 | 0.11 | 0.11 |
| PAX4 | 0.94 | 0.97 | 0.93 | 0.96 | 0.94 | 0.35 | 0.94 | 0.96 | 0.95 | 0.92 | 0.95 | 0.96 | 0.94 | 0.97 | 0.96 | 0.96 | 0.93 | 0.97 | 0.96 | 0.96 | 0.1 | 0.97 | 0.97 |
| PAX6 | 0.65 | 0.1 | 0.63 | 0.1 | 0.11 | 0.49 | 0.69 | 0.85 | 0.48 | 0.41 | 0.1 | 0.1 | 0.48 | 0.49 | 0.49 | 0.49 | 0.11 | 0.1 | 0.88 | 0.48 | 0.13 | 0.49 | 0.48 |
| PDX1 | 0.7 | 0.96 | 0.55 | 0.13 | 0.45 | 0.24 | 0.12 | 0.12 | 0.12 | 0.19 | 0.46 | 0.46 | 0.48 | 0.12 | 0.13 | 0.88 | 0.13 | 0.13 | 0.47 | 0.47 | 0.11 | 0.12 | 0.91 |
| PTF1A | 0.05 | 0.04 | 0.06 | 0.07 | 0.09 | 0.1 | 0.08 | 0.5 | 0.08 | 0.12 | 0.07 | 0.08 | 0.07 | 0.5 | 0.83 | 0.04 | 0.46 | 0.51 | 0.08 | 0.08 | 0.59 | 0.09 | 0.05 |
| RBPJ | 0.26 | 0.23 | 0.26 | 0.24 | 0.57 | 0.35 | 0.26 | 0.01 | 0.25 | 0.84 | 0.23 | 0.01 | 0.24 | 0.01 | 0.01 | 0.24 | 0.87 | 0.01 | 0.01 | 0.01 | 0.76 | 0.01 | 0.01 |
| SHH | 0.04 | 0 | 0.04 | 0 | 0 | 0.17 | 0.03 | 0 | 0.01 | 0.03 | 0 | 0 | 0.01 | 0 | 0 | 0.01 | 0 | 0 | 0 | 0 | 0.07 | 0 | 0 |
| SOX9 | 0.35 | 0.33 | 0.35 | 0.31 | 0.9 | 0.75 | 0.3 | 0.1 | 0.31 | 0.9 | 0.33 | 0.1 | 0.34 | 0.1 | 0.1 | 0.33 | 0.94 | 0.1 | 0.1 | 0.1 | 0.92 | 0.1 | 0.1 |
| TGFB | 0.3 | 0 | 0.31 | 0 | 0.01 | 0.39 | 0.28 | 0.1 | 0.18 | 0.35 | 0 | 0 | 0.21 | 0.06 | 0.06 | 0.2 | 0.01 | 0 | 0.1 | 0.06 | 0.02 | 0.06 | 0.06 |
| TSHZ1 | 0.24 | 0.95 | 0.25 | 0.91 | 0.89 | 0.29 | 0.2 | 0.13 | 0.43 | 0.44 | 0.92 | 0.94 | 0.43 | 0.43 | 0.42 | 0.44 | 0.88 | 0.93 | 0.12 | 0.44 | 0.86 | 0.44 | 0.46 |
| WNT | 0.22 | 0 | 0.22 | 0 | 0.01 | 0.68 | 0.18 | 0.01 | 0.06 | 0.31 | 0 | 0 | 0.08 | 0 | 0 | 0.08 | 0.01 | 0 | 0.01 | 0 | 0.88 | 0 | 0 |

**Table S4 Agreement ratio of comparison between modeling results and scRNA-seq data under various threshold of gene activity.** When the threshold is set to 0.50-0.65, the results (Figure S3) showed the degree of agreement between different steady-state and corresponding cell type expression data in the network calculation. This result is in line with our expectations. Reasons: 1) The network itself does not cover all information; 2) The single cell data itself has an error rate of about 20%. 3). Since the expression of the factors in the steady state is calculated directly through the network, the steady-state characteristics of the high-dimensional nonlinear dynamic system are difficult to manipulate or fit. This result indicates that the partial steady-state correspondence of the network calculation is obtained. Different cell types in mature pancreatic cells.

| Threshold ／ Agreement ratio | 0.5 | 0.55 | 0.60 | 0.65 |
|---|---|---|---|---|
| Alpha cell | 0.500 | 0.577 | 0.653 | 0.731 |
| Beta cell | 0.712 | 0.712 | 0.788 | 0.865 |
| Acinal cell | 0.500 | 0.577 | 0.730 | 0.84 |
| Duct cell | 0.577 | 0.673 | 0.769 | 0.750 |
| Delta cell | 0.635 | 0.558 | 0.827 | 0.865 |

### Supplementary Figures

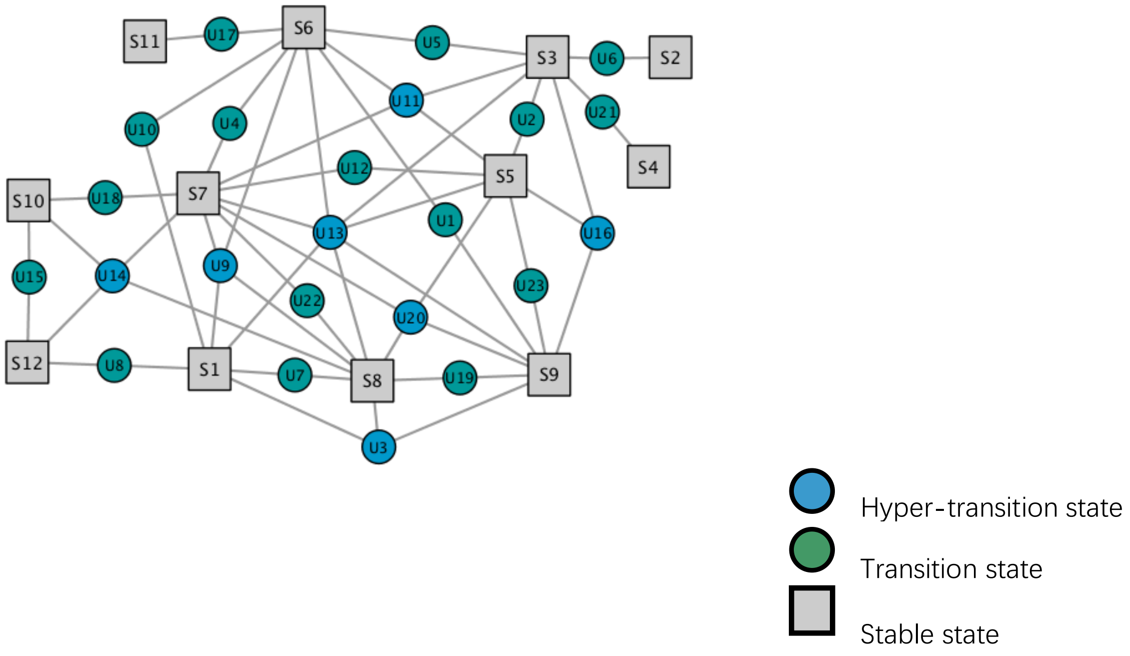


**Figure S1 State connection graph through perturbation analysis describing the possible routes of cell state trans-conversion.** We obtained 12 stable states, 7 hyper-transition states as well as 16 transition states. Through small random perturbation, we obtained the trajectories from each unstable state to it connected stable states and recorded the unstable states that each trajectory passed through, which forms the state connection graph.

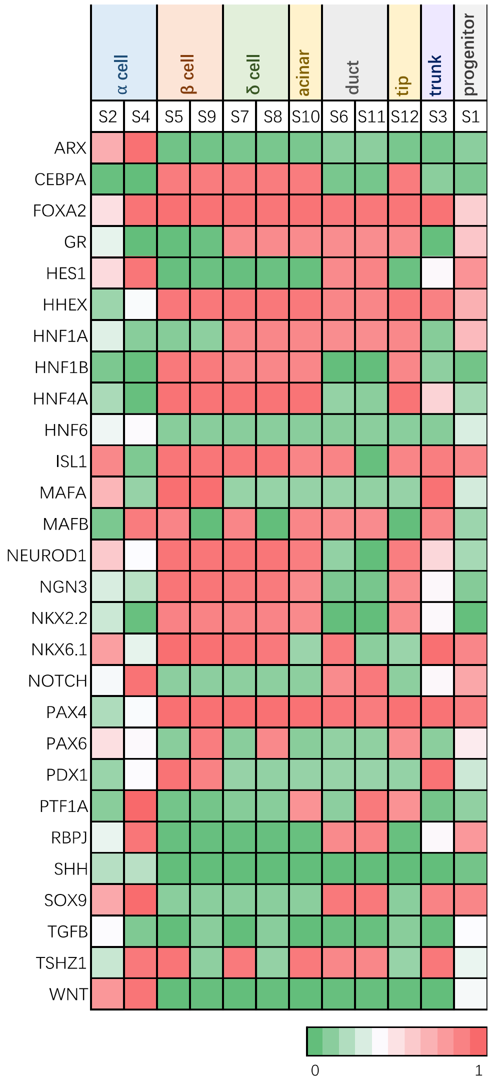


**Figure S2 Linking biological meanings to corresponding stable states.** According to the known marker of specific phenotypic state of cell within pancreas, the biological meanings can be linked to corresponding states.

| | β cell | S5 | S9 |
|---|---|---|---|
| ARX | 0.40 | 0.39 | 0.39 |
| CEBPA | 0.39 | 0.59 | 0.59 |
| FOXA2 | 0.63 | 0.55 | 0.54 |
| GR | 0.44 | 0.25 | 0.26 |
| HES1 | 0.36 | 0.33 | 0.33 |
| HHEX | 0.36 | 0.52 | 0.52 |
| HNF1A | 0.49 | 0.27 | 0.28 |
| HNF1B | 0.36 | 0.60 | 0.60 |
| HNF4A | 0.32 | 0.57 | 0.57 |
| ISL1 | 0.44 | 0.52 | 0.52 |
| MAFA | 0.69 | 0.69 | 0.68 |
| MAFB | 0.57 | 0.61 | 0.30 |
| NEUROD1 | 0.55 | 0.58 | 0.58 |
| NKX2.1 | 0.65 | 0.58 | 0.58 |
| NKX6.1 | 0.69 | 0.55 | 0.55 |
| NOTCH | 0.38 | 0.34 | 0.34 |
| PAX4 | 0.52 | 0.51 | 0.51 |
| PAX6 | 0.56 | 0.31 | 0.71 |
| PDX1 | 0.69 | 0.72 | 0.69 |
| PTF1A | 0.39 | 0.36 | 0.36 |
| RBPJ | 0.39 | 0.34 | 0.34 |
| SHH | 0.39 | 0.39 | 0.40 |
| SOX9 | 0.35 | 0.34 | 0.34 |
| TGFB | 0.36 | 0.34 | 0.44 |
| TSHZ1 | 0.68 | 0.62 | 0.26 |
| WNT | 0.63 | 0.38 | 0.39 |

| | α cell | S4 |
|---|---|---|
| ARX | 0.69 | 0.89 |
| CEBPA | 0.38 | 0.26 |
| FOXA2 | 0.50 | 0.47 |
| GR | 0.33 | 0.25 |
| HES1 | 0.37 | 0.69 |
| HHEX | 0.36 | 0.07 |
| HNF1A | 0.66 | 0.24 |
| HNF1B | 0.37 | 0.26 |
| HNF4A | 0.41 | 0.20 |
| ISL1 | 0.51 | 0.00 |
| MAFA | 0.40 | 0.30 |
| MAFB | 0.63 | 0.60 |
| NEUROD1 | 0.59 | 0.22 |
| NKX2.1 | 0.51 | 0.24 |
| NKX6.1 | 0.41 | 0.18 |
| NOTCH | 0.28 | 0.72 |
| PAX4 | 0.36 | 0.04 |
| PAX6 | 0.62 | 0.33 |
| PDX1 | 0.35 | 0.35 |
| PTF1A | 0.39 | 0.76 |
| RBPJ | 0.38 | 0.70 |
| SHH | 0.39 | 0.56 |
| SOX9 | 0.34 | 0.66 |
| TGFB | 0.35 | 0.36 |
| TSHZ1 | 0.41 | 0.58 |
| WNT | 0.34 | 0.83 |

| | Acinal cell | S10 |
|---|---|---|
| ARX | 0.39 | 0.40 |
| CEBPA | 0.69 | 0.59 |
| FOXA2 | 0.43 | 0.54 |
| GR | 0.68 | 0.59 |
| HES1 | 0.58 | 0.33 |
| HHEX | 0.39 | 0.52 |
| HNF1A | 0.47 | 0.59 |
| HNF1B | 0.52 | 0.58 |
| HNF4A | 0.67 | 0.57 |
| ISL1 | 0.33 | 0.48 |
| MAFA | 0.39 | 0.34 |
| MAFB | 0.34 | 0.61 |
| NEUROD1 | 0.31 | 0.56 |
| NKX2.1 | 0.33 | 0.56 |
| NKX6.1 | 0.38 | 0.20 |
| NOTCH | 0.60 | 0.34 |
| PAX4 | 0.36 | 0.51 |
| PAX6 | 0.33 | 0.31 |
| PDX1 | 0.40 | 0.35 |
| PTF1A | 0.69 | 0.71 |
| RBPJ | 0.69 | 0.34 |
| SHH | 0.39 | 0.39 |
| SOX9 | 0.58 | 0.34 |
| TGFB | 0.55 | 0.34 |
| TSHZ1 | 0.33 | 0.61 |
| WNT | 0.47 | 0.38 |

| | δ cell | S7 | S8 |
|---|---|---|---|
| ARX | 0.39 | 0.40 | 0.40 |
| CEBPA | 0.37 | 0.59 | 0.59 |
| FOXA2 | 0.45 | 0.54 | 0.53 |
| GR | 0.36 | 0.59 | 0.59 |
| HES1 | 0.34 | 0.33 | 0.33 |
| HHEX | 0.68 | 0.52 | 0.52 |
| HNF1A | 0.32 | 0.59 | 0.59 |
| HNF1B | 0.36 | 0.58 | 0.58 |
| HNF4A | 0.37 | 0.57 | 0.57 |
| ISL1 | 0.65 | 0.52 | 0.52 |
| MAFA | 0.40 | 0.34 | 0.34 |
| MAFB | 0.40 | 0.61 | 0.30 |
| NEUROD1 | 0.51 | 0.57 | 0.57 |
| NKX2.1 | 0.44 | 0.57 | 0.57 |
| NKX6.1 | 0.38 | 0.54 | 0.54 |
| NOTCH | 0.45 | 0.34 | 0.34 |
| PAX4 | 0.66 | 0.51 | 0.51 |
| PAX6 | 0.44 | 0.31 | 0.68 |
| PDX1 | 0.46 | 0.35 | 0.35 |
| PTF1A | 0.39 | 0.38 | 0.38 |
| RBPJ | 0.38 | 0.34 | 0.34 |
| SHH | 0.39 | 0.39 | 0.40 |
| SOX9 | 0.37 | 0.34 | 0.34 |
| TGFB | 0.36 | 0.34 | 0.44 |
| TSHZ1 | 0.46 | 0.61 | 0.27 |
| WNT | 0.30 | 0.38 | 0.39 |

| | Duct cell | S6 | S11 |
|---|---|---|---|
| ARX | 0.39 | 0.43 | 0.43 |
| CEBPA | 0.45 | 0.28 | 0.28 |
| FOXA2 | 0.26 | 0.52 | 0.52 |
| GR | 0.46 | 0.58 | 0.58 |
| HES1 | 0.62 | 0.68 | 0.70 |
| HHEX | 0.47 | 0.49 | 0.49 |
| HNF1A | 0.34 | 0.58 | 0.58 |
| HNF1B | 0.66 | 0.26 | 0.26 |
| HNF4A | 0.50 | 0.25 | 0.24 |
| ISL1 | 0.33 | 0.49 | 0.00 |
| MAFA | 0.39 | 0.34 | 0.33 |
| MAFB | 0.34 | 0.60 | 0.60 |
| NEUROD1 | 0.31 | 0.22 | 0.17 |
| NKX2.1 | 0.33 | 0.24 | 0.24 |
| NKX6.1 | 0.41 | 0.53 | 0.19 |
| NOTCH | 0.57 | 0.71 | 0.74 |
| PAX4 | 0.37 | 0.50 | 0.49 |
| PAX6 | 0.32 | 0.32 | 0.32 |
| PDX1 | 0.38 | 0.35 | 0.36 |
| PTF1A | 0.40 | 0.39 | 0.76 |
| RBPJ | 0.43 | 0.70 | 0.71 |
| SHH | 0.69 | 0.39 | 0.40 |
| SOX9 | 0.62 | 0.66 | 0.66 |
| TGFB | 0.64 | 0.36 | 0.35 |
| TSHZ1 | 0.39 | 0.59 | 0.59 |
| WNT | 0.52 | 0.39 | 0.39 |

**Figure S3 Validation of the biological meanings with gene profiling data at molecular level.** Using the normalized methods (See above), the gene activities in stable states from simulation results can be validated and compared with single cell RNA sequencing data. The agreement ratios between the results and independent data are showed in Table S4.
